# Delta integrates 3D physical structure with topology and genomic data of chromosomes

**DOI:** 10.1101/199950

**Authors:** Bixia Tang, Feifei Li, Jing Li, Wenming Zhao, Zhihua Zhang

## Abstract

**Motivation:** The regulation of gene transcription and DNA replication are tightly associated with the 3D chromosomal structures and genomic features, e.g. epigenetic marks, transcription factor bindings and non-coding RNAs. The interaction between the features and the chromosomal structures forming a multilayer 3D regulatory network. Therefore, it is necessary to integrate the physical 3D architecture of genome and features to comprehensive depict their connection to gene regulation.

**Results:** Here, we present an integrative visualization and analysis platform, *Delta*, to facilitate visually annotating and exploring the 3D physical architecture of genomes. *Delta* takes Hi-C or ChIA-PET contact matrix as input and predicts the topology associated domains and chromatin loops in the genome, and generates a physical 3D model which represents the plausible consensus 3D structure of the genome. *Delta* features a highly interactive visualization tool, which enhanced the integration of genome topology/physical structure and extensive genome annotation, by juxtaposition of the 3D model with diverse genomic assay outputs. Finally, we showcased that *Delta* could be helpful to reveal potentially interesting findings by a case study on the β-globin gene region.

**Availability and implementation:** http://delta.big.ac.cn/.

**Supplementary information:** Supplementary data are available at Bioinformatics online.

## 1. Introduction

Many eukaryote cellular processes, e.g. transcription regulation and DNA replication, depend on the spatial architecture of genome in the nucleus (Li, et al., 2012; Roy, et al., 2011; Zhang, et al., 2013). By microscopy-based imaging approaches, large scale architecture of genome has started to be unveiled. For example, each individual chromosome occupies a distinct spatial region, termed chromosome territory, in the interphase nucleus (Hubner and Spector, 2010). In the last decade, the proximity ligation-based chromosome conformation capture (3C) and its variations have substantially improved the resolution and throughput in the exploration of 3D genome architecture (de Wit and de Laat, 2012), and constitute a major engine driving in the field (Bonev and Cavalli, 2016). Upon the date of this paper was writing, the global 3D architecture of the human (Lieberman-Aiden, et al., 2009), mouse (Dixon, et al., 2012) and other mammals (Vietri Rudan, et al., 2015), fly (Sexton, et al., 2012) and yeast genomes (Duan, et al., 2010) have been explored using a high-throughput version of 3C, Hi-C (Lieberman-Aiden, et al., 2009). Mediator-protein-specific 3D chromatin interaction maps have also been surveyed by the ChIA-PET method in mammals for such proteins as CTCF(Handoko, et al., 2011), Pol II (Tang, et al., 2015), cohesin (Demare, et al., 2013) and histone modifications (Heidari, et al., 2014). With these mappings, genomes were found to be physically separated into active and inactive compartments (A and B, respectively) (Lieberman-Aiden, et al., 2009), and further divided into so-called "topologically associating domains" (TADs) (Dixon, et al., 2012; Nora, et al., 2012), and sub-TAD (Phillips-Cremins, et al., 2013; Rao, et al., 2014), along with detailed chromatin looping structures (Rao, et al., 2014; Tang, et al., 2015).

It has been shown that the 3D genome architectures are strongly correlated with genomic features, e.g. histone modifications (Heidari, et al., 2014), DNase sensitivity (Heidari, et al., 2014), transcriptional activity (Tang, et al., 2015), DNA methylation (Wang, et al., 2016) and long noncoding RNA (Quinodoz and Guttman, 2014). Moreover, some of those correlation may be even causal, as the depletion of such elements can result in 3D structure alterations and even diseases (Groschel, et al., 2014; Ibn-Salem, et al., 2014; Lupianez, et al., 2015; Zuin, et al., 2014). Thus, data integration is vital in depicting the complicated interferences of these genomic features and chromatin’s spatial organization. Visualization is always an easy and powerful approach to initial data integration. Many visualization tools have been developed for linear data integrations, such as the genome browsers (Buels, et al., 2016; Stein, et al., 2002; Tyner, et al., 2017). However, to visualize genome features in the 3D genomes, two additional layers of complication emerged as major challenges. First, the topology of correlations between genomic loci forms a nonlinear complex network. As distal genome loci may physically contact to each other via chromatin looping in 3D space, the genomic features may highly correlated at the interacting loci. However, it is not fully supported to visualize the topology of this complex network in the traditional genome browsers. Second, limited by physical space, it is nearly unavoidable to have interfering genome features when the number of features that to be added into a physical 3D model increases. Several visualization tools have been developed for 3D genome data (Durand, et al., 2016; Lajoie, et al., 2009; Li, et al., 2016; Mohamed Nadhir Djekidel, et al., 2016; Paulsen, et al., 2014; Xu, et al., 2016; Yanli Wang BZ, 2017; Zhou, et al., 2013), and some of the tools implemented features aimed to the above challenges in one or another aspects, however, the data integration in topology and 3D physical space remains challenging.

To meet these challenges, we developed an easy-to-use web based 3D genome visualization platform, *Delta*, to integrative display 3D structure of genome with diverse genomic features by juxtaposition of the 3D model with nearly unlimited number of genomic assay outputs. This unique feature distinguishes Delta from current visualization tools. Moreover, we showcased *Delta* by applying it to study a well-studied locus control region (LCR) using public data. By visually inspecting the paradoxical 3D physical model of LCR-β-globin gene region and the multi-omics data, we speculated that Hi-C loop called between the 5th 5’ DNase I hypersensitive site (5’HS5) and 3’ HS1 may indicate a constructive and transient interaction between the loci. On the other hand, while internal four 5’ HSs have closer interaction with beta-globin genes, the interaction may be stochastic, and open chromatin structures are essential toward the expression of beta-globin genes. This speculation is further evidenced by literature search, suggesting that Delta could be helpful to infer novel and valid hypothesis via visually integrating multiple datasets.

## 2. System and methods

### 2.1 Basic usage of Delta

Delta is composed with *Delta-analysis*, an easy-to-use pipeline to process user provided Hi-C/ChIA-PET data, and *Delta-view*, an integrative visualization tool. The Delta-analysis is an assembly of tools to perform TAD calling (Weinreb and Raphael, 2016), chromatin loop calling (Xu, et al., 2016) and physical 3D modeling from 3D genome data (Hu, et al., 2013; Trieu and Cheng, 2016). Delta-analysis take Hi-C or ChIA-PET contact matrix as input (Dixon, et al., 2012; Lieberman-Aiden, et al., 2009; Nora, et al., 2012; Rao, et al., 2014; Tang, et al., 2015), and results can be directly dumped into the Delta-view. Alternatively, the users can directly upload pre-calculated TAD, chromatin loop and/or physical 3D model to Delta-view. The Delta-view can be run under either uni-mode or dual-mode. Under the uni-mode, the users can choose one of the following three modes, linear genome mode, circlet mode and 3D physical mode (Figure 1). Compared to the JBrowser (Buels, et al., 2016), the linear genome mode in Delta-view can display more track types, such as contact matrix as 45-degree rotated half heatmap, to facilitate better aligning with additional linear genomic features in the linear view. Delta-view allows users to adding tracks in the 3D physical model to probe the interested genome context in detail. Under the dual-mode, Delta-view makes a juxtaposition of 3D physical model with a conjunctive genome browser or circlet viewer (see section 2.5 for details).

**Figure 1.**
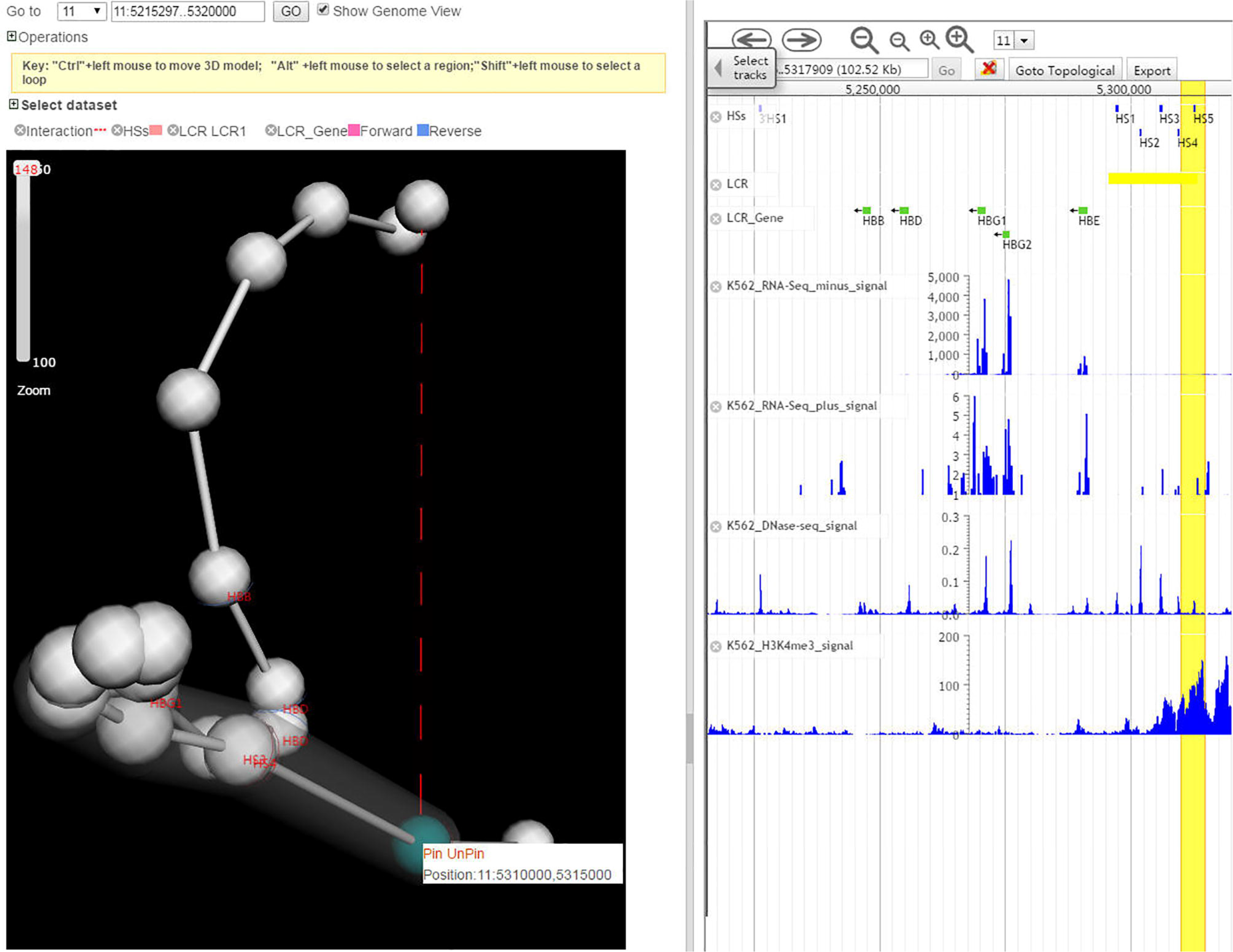
Screen shots of three basic mode of Delta. A) Genome view, B) circlet view and C) 3D physical view.

### 2.2 Visualizing 3D physical model with genomic features

Delta-view presents the 3D physical structure of a given chromosome (fragment) with ball-and-stick model. Delta-view’s 3D physical mode takes a coordinate file as input. The coordinate file can be provided either by users or generated by Delta-analysis from contact matrix. In the ball-and-stick model, each bead represents a bin which is a user-defined genome region as a basic unit while generating the coordinate file. In the present version of Delta-view, the bin size is given by the input contact matrix file, zooming-in/out does not change the resolution of the 3D physical model *per se.* However, the zooming-in/out may indeed alter the resolution of annotation tracks which are printed on the surface of the beads. In another word, Delta-view can automatically determine the way how a genomic feature can be displayed at the current zooming status, to maximize the readable information displaying.

### 2.3 Adding tracks onto the 3D physical model

Tracks, as it was defined in genome browsers, are individual plots that represent genomic annotation data (features)(Buels, et al., 2016; Kent, et al., 2002; Skinner, et al., 2009). Incorporating genomic features into 3D physical model is helpful for assaying the association of the two. Due to possible interference between tracks in the limited space on the ball-and-stick model, Delta-view restrict the number of maximal tracks could be displayed in a 3D physical model to 5. Four types of tracks (quantitative, regional, labeling, and connective) are currently supported (Figure 2).

**Figure 2.**
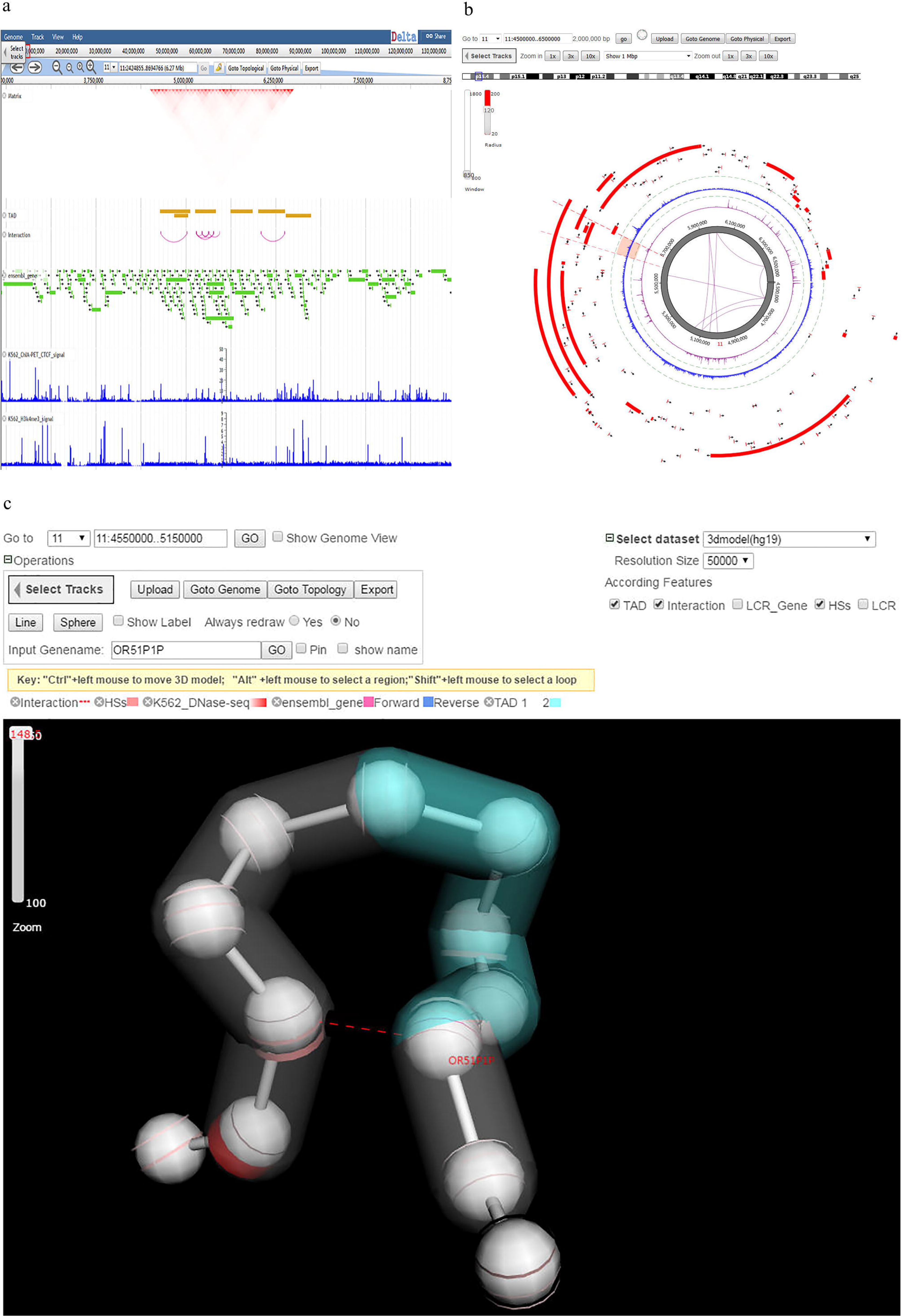
The examples of four type of annotated tracks in the 3D physical model. The model was generated from Hi-C data in K562 cells (chr11, from 4,600,000 to 5,250,000) with resolution at 50k binsize. A). Quantitative feature. The colored stripes represent the DNase-seq reads density in the region. B). Regional feature. The shadows represent the TADs called from Hi-C data. C). Labeling feature. Several the genes in the region are labeled by their name. D) Connective feature. The dashed arcs represent chromatin loops called from Hi-C data in the region.

The quantitative feature, as the major category of features that covers the output of a wide spectrum of biochemical assays, e.g. ChIP-seq, DNase-seq, RNA-seq, is represented as a colored stripe on the beads (Figure 2A). The brightness and width of the stripe represents the strength of the feature. Delta-view can automatically determine to display summarized data or raw data points according to the current zooming level. For example, when the mode is fully zoomed out, the mean value of the feature within the genome region of a bead is painted onto the whole bead, while zooming in, more and more detailed features will emerge. Ideally, i.e the model was built with sufficient small bin size, each annotated ChIP-seq peaks shall be presented when zoom into the highest resolution. However, because the highest resolution 3D physical models that possibly generated from current Hi-C data is approximately a hundred kilobases, we only allow maximal 10 bins within a bead for any quantitative features.

The regional feature is designed to visualize the large genomic domains (Figure 2B). It has been recognized that genome is organized into hierarchical domain structure, such as gene-rich domains (Gilbert, et al., 2004), or TADs (Dixon, et al., 2012; Nora, et al., 2012). It has been suggested that the TAD may be resulted from hierarchical folding dynamic of genome (Dixon, et al., 2012; Nora, et al., 2012), and has been proven strongly associated with functional genomic or epigenomic domains (Huang, et al., 2015). Therefore, it is vital for learning the domain structure comprehensively to visualize the domain in 3D physical model together with quantitative features. A regional feature is visualized as a colored shadow covering the beads within the domain. The minimal unit in displaying regional feature is a bead, in another word, when a domain cover a region less than a bead represented, the whole bead will be shadowed. If there are more than two domains need to be shown in the current visual field, Delta-view will shadow the domains with two alternating colors. Users can upload their customized domain data as well.

The labeling feature is represented as plain text marked next to the genomic features (Figure 2C). The most commonly used labeling feature is gene name. Although having all genes labelled is supported, it is not recommended as the labels will be overlapping with each other and can be hardly distinguished. Users can label the selected genes by inputting the names. In fact, Delta-view takes the gene annotations as both quantitative and labeling features. When the view is extremely zoomed out, Delta-view will take the genes as quantitative features with brightness of color represents the gene density in the genome region, when the view is sufficiently zoomed in, the gene features will be turned into labeling, in which each gene can be presented as individual stripe with gene name labeled, the orientation of the gene is indicated by the color of stripe.

The connective features is the dashed arcs linking two beads (Figure 2D), representing the chromatin loops (Rao, et al., 2014; Tang, et al., 2015; Xu, et al., 2016). Delta-view displays intra-region (visual field) loops, i.e. the loops connecting to the beads out of current visual field will be omitted. This setting can help users focus on the selected partners in a clear vision. The arcs can be highlighted using mouse click with “shift” key down. Highlighting an arc will refresh the conjunct genome/circlet browser (see below). This is rather useful when comparing the two connected loci with multiple epigenetic features.

### 2.4 Pinning a bead in a 3D physical model

It is vital to rotate a physical model. While rotating the model, it is quite easy to get lost if there are too many components in the visual field. Hence, in Delta-view users can pin a bead at the center of the visual field, i.e. mimicking an invisible pin anchored a selected bead at the center of the visual field and the pinned bead is always stay at the center no matter how users rotate the model. Besides the beads, the genes can also be pinned while labeling. Labelling gene names and pinning may facilitate to explore the gene specific or enhancer specific promoter-enhancer interactions.

### 2.5 Dual-mode with linear and circlet view

Under the dual-mode, a 3D physical model and a conjunctive view are juxtaposed in Delta-view (Figure 3). To enter the dual-mode, users can click on the check box named “Genome view” in the 3D physical view page. Once entering into the dual-mode, the data displayed in the conjunctive linear or circlet view is always synchronized with the physical view. For example, if you select a part of the physical model by drawing mouse left key with “Alt” key down, the conjunctive linear or circlet view will zoom into the given region automatically. When users highlight an object in the physical view, e.g. a bead or an arc, the corresponding region will also be highlighted in the conjunctive view immediately.

**Figure 3.**
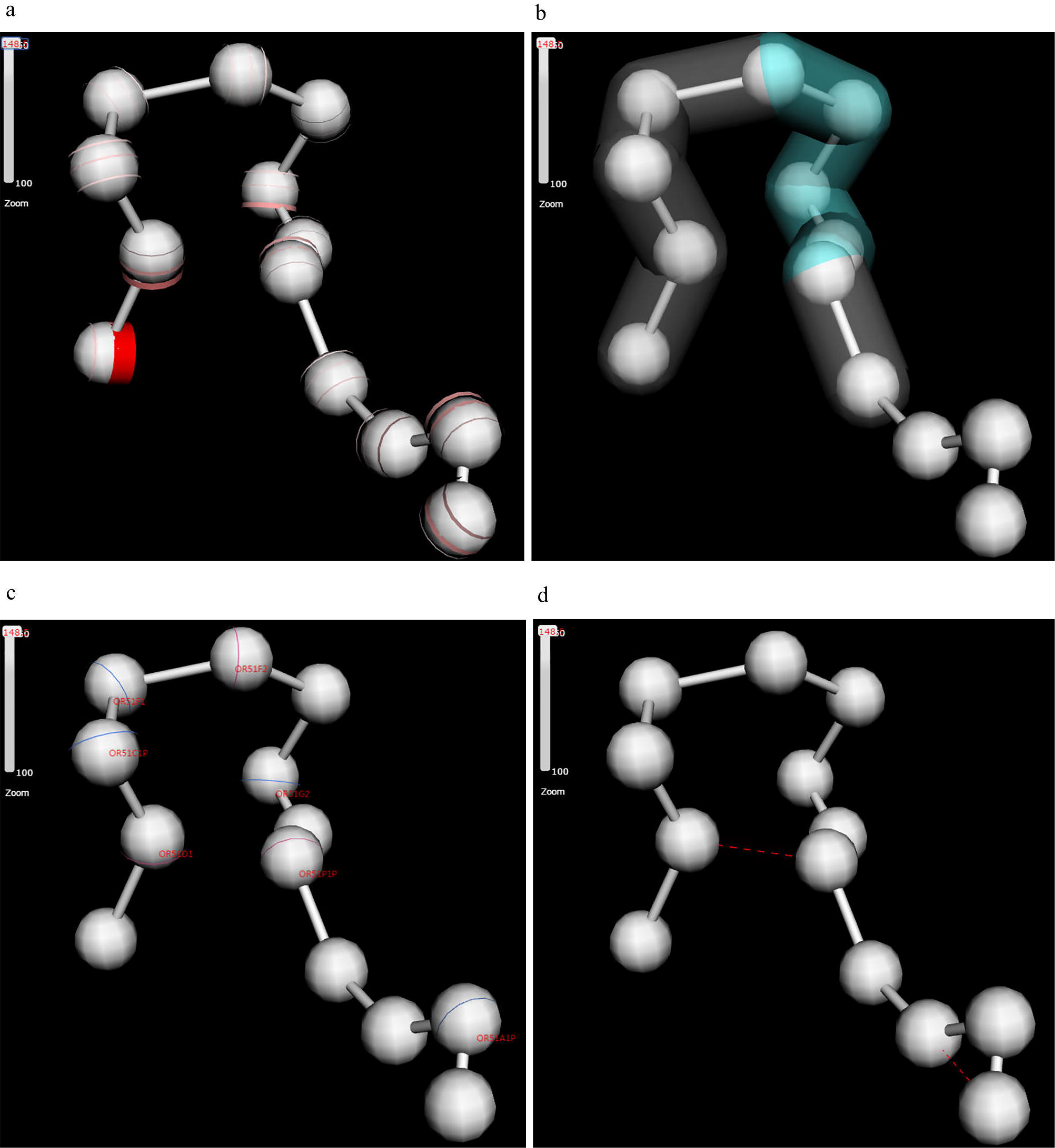
The β-globin locus region shown in the dual mode. The model was cropped from a larger model as in Figure 2 with resolution at 5k binsize (chr11, from 5,215,297 to 5,320,000). Because the LCRs is defined as regional features, it is shadowed in grey. The HSs and β-globin genes are labeled with their names. In the physical model, the bead where 5’HS5 located in is pinned and highlighted, the corresponding region is also highlighted in the conjunctive genome browser window. Multiple genomic data, including RNA-seq, DNase-seq, ChIP-seq of H3K4me3 and gene annotations are shown in the conjunctive genome browser.

The dual-mode let Delta-view overcomes the limitation of maximal tracks that is allowed to be simultaneously displayed in the 3D physical model, as one can add unlimited tracks in the conjunctive views. Because of this restriction, the newly added tracks in physical model is always synchronized to the conjunctive view immediately, while newly added tracks in the conjunctive view are not immediately updated to the physical view. In this case, the users have to manually click the “update physical model” button, which will trigger a pop-up dialog box asking the selection of tracks that need to be synchronized when there are more than five tracks active in the conjunctive view.

## 3. Implementation

Delta is developed based on J2EE framework with MySQL as its database engine. The user interface is developed by JavaScript, HTML, and AJAX. The linear genome view was a modified version of JBrowse (Buels, et al., 2016; Skinner, et al., 2009). The circle mode was modified from Epigenome browser (Zhou, et al., 2013). The physical view was developed based on 3Dmol.js (Rego and Koes, 2015). The Delta-analysis takes a contact matrix as input. The associated meta-data, such as organism name, bin size and the targeted genome region, are required while uploading the input data. The TADs, the chromatin interaction loops and the 3D physical model were calculated by TADtree (Weinreb and Raphael, 2016), FastHiC (Xu, et al., 2016), and BACH (Hu, et al., 2013) and MOGEN (Trieu and Cheng, 2016), respectively. Each session has a unique session number which can be used to retrieve the processed data.

The pre-installed Hi-C datasets were from publications (Dixon, et al., 2012; Lieberman-Aiden, et al., 2009; Rao, et al., 2014), genomic features were downloaded from ENCODE project (Consortium, 2013), genome annotation were downloaded from Ensembl (Kinsella, et al., 2011). It shall be noted that the data we downloaded were processed peaks which were called by the origin data centers, thus two reference genomes assemble (hg18 and hg19) were included. For circlet and physical mode, both of them take the following data procedure, first, the bigwig signal files were converted to bedGraph format using Kent source utilities bigWigToBedGraph executable (Kent, et al., 2010), then, a Tabix indexed files were generated from the bedGraph file to determine the zoom scale. The detail information of used data can be found from http://delta.big.ac.cn/circosweb/pages/dataset/dataset.jsp.

### 3.1 Availability of data and material

The Delta is made open source at (https://github.com/zhangzhwlab/delta) and accessible from http://delta.big.ac.cn. A standalone version of Delta is also available from http://delta.big.ac.cn/pages/download/download.jsp. For more information about the Delta please visit the help page in the website.

## 4. Discussion

### 4.1 Transient interaction between constructive chromatin loop in the LCR

To illustrate the potential usefulness of the Delta for interesting biology insights, we focused on a canonical β-globin locus, which has a complex structure and developmental gene activation pattern (Palstra, et al., 2003). The human β-globin locus contains a family of genes that are activated by a distal enhancer cluster, the locus control region (LCR), that is characterized by five DNase I hypersensitive sites (HSs) (Figure 3). To better understand the regulatory network of LCR-β-globin locus, we modeled the physical 3D structure of this region and put it in a juxtaposition with genome browser (Figure 3). The contact matrix data were downloaded from the high resolution in-situ Hi-C data (Rao, et al., 2014), and the region we chose for modeling is larger (Chr11: 4,500,000-6,500,000) than 5’HS5-3’HS1 region as the local structure may influenced by external sequences. The 3D physical models at two resolutions (50kb and 5kb bin size) were generated by Delta-analysis. We noticed that the LCR indeed closely attached to the globin genes in the physical 3D model with none special loop were identified between any particular HSs and globin genes. If the LCR contact is sufficient to the expression of the globin genes, we would expect to see the active expression of all globin genes in K562 cells. However, RNA-seq data indicates that only HBE, HBG1 and HBG2 gene are actively expressed. Thus, we lineup DHS and H3K4me3 ChIP-seq data and found that indeed the promoter regions of three genes are more accessible as well as enriched for actively promoter marks. In the promoter regions of HBD and HBB genes, although the DNase-seq data showed a moderate peak, there were no obvious spotted peaks of H3K4me3. This visualization pattern implies that the contacts from LCR, accessible promoter with active histone mark may be necessary to the active expression of globin genes, however, none of the three may be sufficient. We also observed that 5’HS5-3’ HS1 chromatin loop which called from Hi-C data is not connecting the spatially approximated beads in our physical 3D model (Figure 3). This observation is less likely due to computational artifacts as it has been revealed in both resolutions of physical 3D models from multiple Hi-C and ChIA-PET datasets (Figure S1). Because the physical 3D model was a consensus average from a structure ensemble (Hu, et al., 2013), we speculated that the loop between 3’HS1 and 5’HS5 may indicate a higher than expect frequency transitorily interactions. This transitorily interactions therefore may function as the gate keeper for LCR-globin looping domain.

To validate this working hypothesis of LCR-β-globin regulatory network, we searched the literatures and found that the transcriptionally active globin genes are indeed closely positioned to the LCR HSs (Palstra, et al., 2003), while this chromatin loop structure is not maintained by the transcription activity of the globin genes *per se*(Mitchell and Fraser, 2008). On the other hand, once the loop between 5’ HS5 and 3’ HS1 were disrupted by CTCF knockdown in K562 cells, the transcription of globin genes was also reduced (Hou, et al., 2010; Palstra, et al., 2003). This example demonstrates the usefulness of visualizing the 3D physical models with various genomic data together to initiation a working hypothesis on a gene regulatory system.

### 4.2 Comparison with other tools

To assess the performance of Delta, we compared it to eight published 3D gnome visualization tools, including five popular visualization tools (Hi-Browse (Paulsen, et al., 2014), Juicebox(Durand, et al., 2016), my5C(Lajoie, et al., 2009), 3D genome browser (Yanli Wang BZ, 2017) and Epigenome browser(Zhou, et al., 2013)), which have been recently reviewed(Yardimci and Noble, 2017), HiCPlotter(Akdemir and Chin, 2015) and HiC-3Dviewer (Mohamed Nadhir Djekidel, et al., 2016) and 3Disease Browser (Li, et al., 2016) (Table 1). The last two also have implemented physical mode. Among all above eight publicly-available tools, Delta is the only one that provides all the linear, circlet and 3D physical visualization modes. Nearly all features in the three modes were implemented in Delta, except for normal heatmap and virtual 4C plot. It’s important to equip dual mode, which juxtaposes two distinct plots in the same window. In Delta, we focused on 3D physical mode with embedded physical-genome and physical-circlet dual modes, while the 3D genome browser and the Epigenome browser have implemented Heatmap-Genome dual-mode and HiC-3Dviewer implemented Heatmap-physical mode. In those dual-modes, the data will be synchronized during the session of visualization and analysis. Epigenome Broswer has the largest pre-loaded datasets and supplemental datasets, followed by Juicebox. Delta has also preloaded most popular publicly-available Hi-C datasets, and users can also either upload local customized datasets or external datasets by provide remote URLs. Taken together, Delta is a competitive visualization tool in the majority aspects of 3D genome to current tools, with a unique merit of interactively and lively dual visualization mode of 3D physical model with genome and circlet plots.

### 4.3 Conclusions

Chromosome architecture in the context of genomic features is critical to understand the function of the genome. Given the complicated nature of the metazoan genome, it is essential to understand the principles behind the nuclear architecture from various perspectives, such as physical 3D structure, topological architecture and linear genomic arrangements. Comprehensively exploring the principles is possible with the advent of technologies, such as imaging, nextgeneration sequencing combined with conformation capture methods, as well as theoretic researches on macromolecule 3D structure and complicated network dynamics. With the launching of NIH’s 4D Nucleome projects, massive sequencing and imaging data is expected to bloom up in the coming years. Thus, the need for an integrative analysis and visualization platform like Delta is urgent. We believe the interactive and integrative feature of Delta will help researchers to initiate meaningful intuition and direct profound associations between datasets. Delta will keep upgraded and extended to new media, such as mobile device in the near further.

## Acknowledgements

This work was supported by grants from the National Nature Science Foundation of China [91540114, 31671342 to ZZ], the National Basic Research (973) Program of China [2014CB542002 to ZZ] and the National High Technology Development 863 Program of China [2014AA021103 to ZZ], this work was also supported by the National Nature Science Foundation of China [31401113, to FL, 31401112 to LJ]. The funders had no role in study design, data collection and analysis, decision to publish, or preparation of the manuscript. We thank Data Center of Beijing Institute of Genomics (BIGD) hosts the Delta and provides computing resources.

## Abbreviations

3D: three dimensional
3C: chromatin conformation capture
4C: circular chromosome conformation capture
5C: chromosome conformation capture carbon copy
TAD: topologically associating domain
ENCODE: Encyclopedia of DNA Elements
LCR: loucs control region

## Author’s contributions

TB contributed code. ZW provided advice on the user interface design. ZZ conceived of and led the project. TB, FL, JL and ZZ wrote the paper. All authors read and approve the final manuscript.

---

Table 1. Comparison of 3D genome visualization tools.

Figure S1. The β-globin locus region shown in the dual mode. The model was cropped from a larger model as in Figure 2 with resolution at 50k binsize (chr11, from 5,068,001 to 5,466,000). Because the LCRs is defined as regional features, it is shadowed in grey. The HSs and β-globin genes are labeled with their names. In the physical model, the bead where 5’HS5 located in is pinned and highlighted, the corresponding region is also highlighted in the conjunctive genome browser window. Multiple genomic data, including RNA-seq, DNase-seq, ChIP-seq of H3K4me3 and gene annotations are shown in the conjunctive genome browser.

